# Mavacamten facilitates myosin head ON-to-OFF transitions and shortens thin filament length in relaxed skeletal muscle

**DOI:** 10.1101/2024.11.29.626031

**Authors:** Michel N. Kuehn, Nichlas M. Engels, Devin L. Nissen, Johanna K. Freundt, Weikang Ma, Thomas C. Irving, Wolfgang A. Linke, Anthony L. Hessel

## Abstract

The first-in-its-class cardiac drug mavacamten reduces the proportion of so-called ON-state myosin heads in relaxed sarcomeres, altering contraction performance. However, mavacamten is not completely specific to cardiac myosin and can also affect skeletal muscle myosin, an important consideration since mavacamten is administered orally and so will also be present in skeletal tissue. Here, we studied the effect of mavacamten on skeletal muscle structure using small-angle X-ray diffraction. Mavacamten treatment reduced the proportion of ON myosin heads but did not eliminate the molecular underpinnings of length-dependent activation, demonstrating similar effects to those observed in cardiac muscle. These findings provide valuable insights for the potential use of mavacamten as a tool to study muscle contraction across striated muscle.

## Introduction

In the presence of calcium (pCa < ∼7), active tension generation in sarcomeres is produced by crossbridge cycling between the myosin motors in the thick filament and actin in the thin filaments (Fig. 1A) (1). The performance of sarcomeres during contraction is partially set in the relaxed condition by manipulating the propensity of myosin heads to form crossbridges upon activation. Slight changes to the number of available myosin heads in relaxed sarcomeres have outsized consequences for contraction (2, 3) and create a unique structural fingerprint that is measurable via small-angle X-ray diffraction (4). In resting sarcomeres (Fig. 1A), myosin heads are in a structural state between an OFF state, where the myosin heads are folded onto their tails and docked into helical tracks along the thick filament backbone, and an ON state, where myosin heads are positioned up and away from the thick filament and towards the thin filament (4, 5). Of note, the sarcomere length-dependence of ON myosins relates to calcium-sensitivity during contraction (6– 8) and underpins the Frank-Starling effect (4, 5, 9). Recently, a first-in-its-class cardiac drug was approved as a myosin motor inhibitor, called mavacamten (marketed as Camzyos) (10), that shifts cardiac myosin motors from the ON toward the OFF state and reduces contraction tension (11– 13). Mavacamten is not specific to the cardiac myosin isoform and also can bind to skeletal muscle, albeit with reduced affinity. In skeletal muscle, the structural effect of mavacamten is unknown but it does reduce active tension with less potency (14) and is more effective in slow vs. fast fiber types (10). We evaluated the myofilament structural signature of resting wildtype skeletal fibers before and after treatment with mavacamten. We found that mavacamten treatment shifts myofilament structures towards an OFF conformation and alters the thin filament in a way consistent with those seen in cardiac sarcomeres, demonstrating mavacamten functionality across striated muscles.

**Figure 1.**
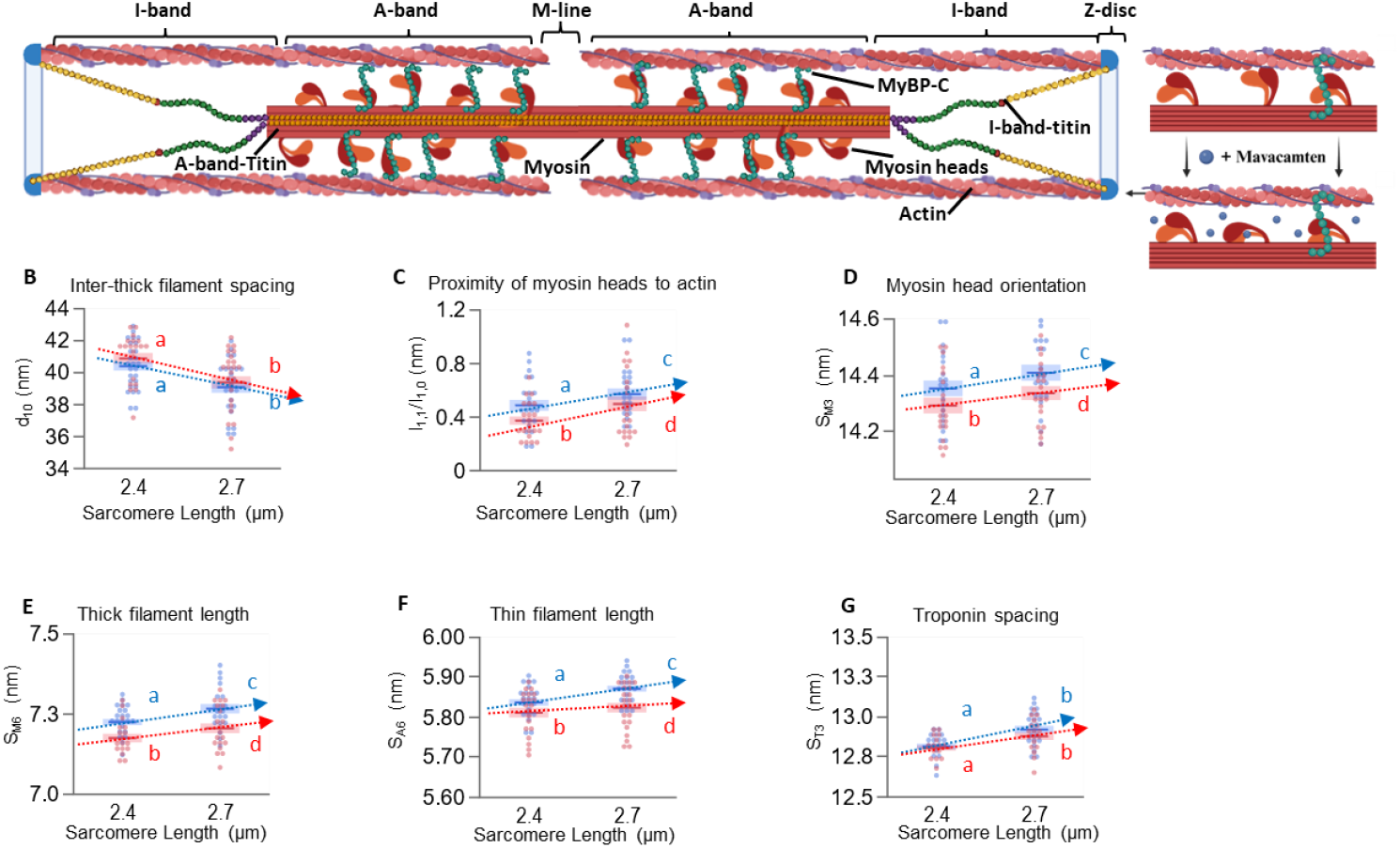
Sarcomeric structures before (blue) and after (red) mavacamten. (A) Cartoon of a passive sarcomere incubated with mavacamten. d_10_ (B), I_1,1/I1,0_ (C), S_M3_ (D), S_M6_ (E), S_A6_ (F) and S_T3_ (G) provided. Data presented as mean ± s.e.m. Statistical significance indicated by connecting letters, where conditions with different letters are significantly different. 2-way ANOVA with individual random effect and repeated measure design. Further details in Table 1.

## Results

X-ray diffraction patterns were collected and analyzed for relaxed permeabilized fiber bundles at 2.4 and 2.7 µm sarcomere length (SL), before and after incubation in mavacamten. Full statistics are in Table 1. Interfilament lattice spacing was quantified as the spacing between the thick-filament containing d1,0 planes of the filament lattice (Fig. 1; Table 1) and decreased from short to long SL, as expected, with no detectable effect of mavacamten (Fig. 1B). Myosin head ON/OFF level was quantified by tracking the ratio of the intensities of the 1,1 and 1,0 reflections (I_1,1_/I_1,0_; Fig. 1C), a measure of relative shifts of mass (e.g., myosin) between the thick and thin filaments, and the spacing of the M3 reflection (S_M3_; Fig. 1D), a measure of the degree of myosin head orientation, with both values positively related to the transition of myosin heads towards the ON state. Before and after mavacamten treatment, I_1,1_/I_1,0_ and S_M3_ increased from short to long SL, indicative of the well-known SL-dependent transition of myosin heads from the OFF towards ON state (4). After mavacamten treatment, I_1,1_/I_1,0_ and S_M3_ decreased across SLs, which suggests an ON-to-OFF transition of the myosin heads. The spacing of the M6 reflection (S_M6_; Fig. 1E) tracks other thick filament activation markers. S_M6_ increased from short to long SL before and after mavacamten treatment, however mavacamten treatment reduced S_M6_ across SLs (Fig 1E; Table 1). Thin filament length was quantified via the spacing of the A6 reflection (S_A6_; Fig. 1F), which represents the periodicity from the pitch of the left-handed axial turn of the actin double helix (4). Before and after mavacamten treatment, S_A6_ increased from short to long SL (Fig. 1F; Table 1). However, mavacamten treatment reduced the S_A6_ across SLs, while not changing the relative magnitude of thin filament elongation from the short to long SL (ANCOVA interaction p = 0.12). Finally, we obtained the periodicity of the troponin structure by the spacing of the T3 reflection (S_T3_; Fig. 1G). S_T3_ increased with stretch, but mavacamten treatment had no significant effect (Fig. 1G; Table 1).

**Table 1.**
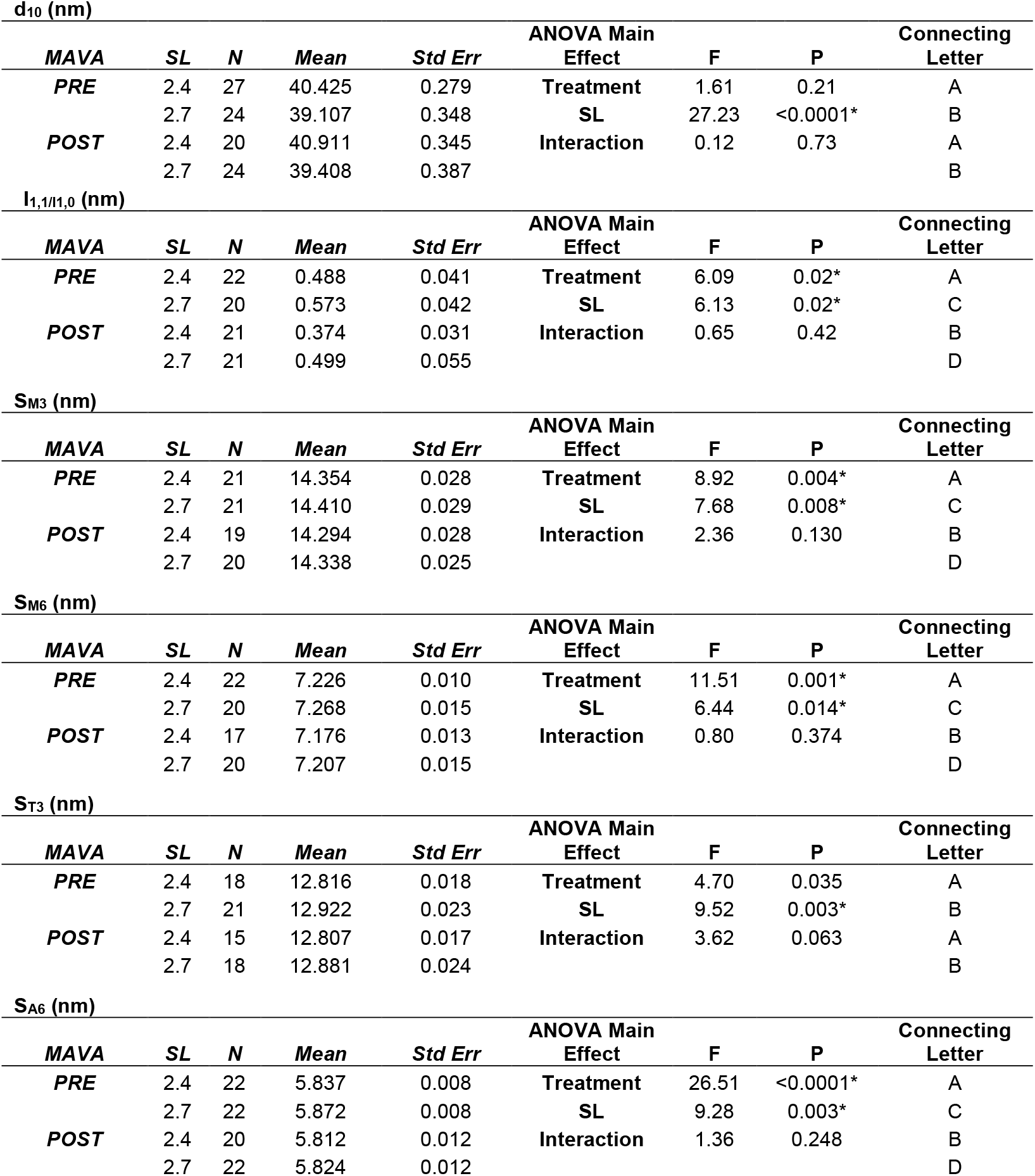
Statistical details from experiments shown in Fig. 1 (B-G). N = number of preparations included in the condition. The ANOVA analysis F-statistics (F) and P-values (P) are provided, as well as a connecting letter report from a Tukey’s honest significant difference (HSD) analysis on sarcomere length (SL) before (PRE) and after (POST) mavacamten (MAVA) treatment. Data reported as mean ± s.e.m. *Significant (P < 0.05).

## Discussion

We demonstrate in skeletal muscle that mavacamten treatment leads to a change in X-ray diffraction signatures that are consistent with a transition of a proportion of myosin heads from the ON to the OFF state (15), similar to that seen in cardiac muscle. In mammals, the control of myosin head configuration is typically regulated by both SL-dependent and SL-independent mechanisms, which is structurally evaluated here by changes in myosin-head associated (e.g. S_M3_ and I_1,1/I1,0_) and thick-filament associated (e.g. S_M6_) X-ray reflections. Physiologically, these control mechanisms are regulated by sarcomeric proteins such as titin and MyBP-C (16, 17). In our study, we over-rode these control schemes with mavacamten, which directly targets the myosin motors and drives them towards an OFF conformation (12, 18). This led to a SL-independent decrease in S_M3,_ I_1,1/I1,0_, and S_M6_, which is a signature of an ON-to-OFF transition of myosin heads (15).

The control of SL-dependent myosin ON/OFF states are thought to underlie myofilament length-dependent activation, the cellular mechanism behind the Frank-Starling Law of the heart, which enhances calcium sensitivity at longer vs. Shorter SLs (8). As sarcomeres stretch, there is a myosin heads OFF-to-ON transition, increasing force production upon activation, likely regulated by the titin I-band spring (16). We observed an OFF-to-ON length-dependent transition of myosin heads regardless of mavacamten treatment. These observations suggest that the intrinsic mechanisms governing length-dependent activation are intact even with the pharmacological suppression of myosin activity and aligns with studies utilizing cardiac tissue (13).

Of note, compared to the short SL, at long SL we identified a positive relationship between increasing myosins in the ON state, and increasing thin filament length (inferred from S_A6_), and this relationship was upheld after mavacamten treatment. In cardiac and skeletal muscle, thin filament proteins can be altered by changing the myosin ON/OFF state balance, suggesting myosin head interactions with actin (6, 8). Crossbridge-based force in passive muscle could stretch and hold the thin filaments. In fact, careful assessment of passive cardiac muscle has made clear that crossbridge tension is present, even if only a few crossbridges are forming (19, 20). Isolating the effects of these crossbridges is challenging due to complex interactions between sarcomeric proteins, therefore computational models are needed to clarify these dynamics.

## Materials and Methods

Animal procedures were approved by LANUV NRW (81-02.04.2019.A472). We prepared permeabilized fiber bundles from mouse psoas muscle (age range 4-8 months old, n = 28 fiber bundles, N = 8 mice [3 male / 5 female]) and placed them into an experimental apparatus in the small-angle diffraction instrument at the BioCAT beamline 18D (Advanced Photon Source; Argonne National Laboratory, USA; (21)) as previously described (6). Fiber bundles (15–30 fibers, 3–6 mm long) were prepared and attached to the experimental rig, as described previously (6). Bundles were subjected to a passive ramp stretch at 0.1 μm SL/s from 2.4 to 2.7 μm SL, before and after a 20 min incubation with 50 µM mavacamten (Fischer Scientific). X-ray exposure up to a total exposure of 1s occurred at steady-state force at both SLs. X-ray diffraction patterns were analyzed using the free software MuscleX (22). Box-Cox transformations were applied, and significant effects were analyzed with a post-hoc Tukey’s highly significantly different multiple comparison procedure. Data presented as mean ± s.e.m.

## Data and materials availability

All data are available in the main text or the supplementary materials, or available upon reasonable request.

## Acknowledgments

Funding for this study was provided by the German Research Foundation (454867250 [ALH], SFB1002A08 [WAL]), IZKF Münster (Li1/029/20 [WAL]), and the National Institute of Health (P41 GM103622 [TCI]), P30 GM138395 [TCI], R01HL171657 [W.M]). BioRender.com was used to construct the figure cartoon.

